# Mutations in the albinism gene *oca2* alter vision-dependent prey capture behavior in the Mexican tetra

**DOI:** 10.1101/2024.06.17.599419

**Authors:** Stefan Choy, Sunishka Thakur, Ellen Polyakov, Jennah Abdelaziz, Evan Lloyd, Maya Enriquez, Nikita Jayan, Yaouen Fily, Suzanne McGaugh, Alex C Keene, Johanna E Kowalko

## Abstract

Understanding the phenotypic consequences of naturally occurring genetic changes, as well as their impact on fitness, is fundamental to understanding how organisms adapt to an environment. This is critical when genetic variants have pleiotropic effects, as determining how each phenotype impacted by a gene contributes to fitness is essential to understand how and why traits have evolved. A striking example of a pleiotropic gene contributing to trait evolution is the *oca2* gene, coding mutations in which underlie albinism and reductions of sleep in the blind Mexican cavefish, *Astyanax mexicanus*. Here, we characterize the effects of mutations in the *oca2* gene on larval prey capture. We find that when conspecific surface fish with engineered mutations in the *oca2* allele are hunting, they use cave-like, wide angle strikes to capture prey. However, unlike cavefish or surface fish in the dark, which rely on lateral line mediated hunting, *oca2* mutant surface fish use vision when striking at prey from wide angles. Finally, we find that while *oca2* mutant surface fish do not outcompete pigmented surface siblings in the dark, pigmented fish outcompete albino fish in the light. This raises the possibility that albinism is detrimental to larval feeding in a surface-like lighted environment, but does not have negative consequences for fish in cave-like, dark environments. Together, these results demonstrate that *oca2* plays a role in larval feeding behavior in *A. mexicanus*. Further, they expand our understanding of the pleiotropic phenotypic consequences of *oca2* in cavefish evolution.

## 1. Introduction

Identifying the genes and genetic changes that underlie trait evolution is central to understanding how and why traits evolve. It is widely recognized that evolution utilizes only a subset of the genes that underlie traits (Conte et al., 2012; Martin and Orgogozo, 2013; Stern, 2013). However, why some genes are repeatedly used by evolution while others are not is not fully understood (Bolnick et al., 2018). Pleiotropy, or the phenomenon of a single genetic locus impacting two or more phenotypic traits, has been proposed as one reason why evolution utilizes some genes in favor of others (Fisher, 1930; Wright, 1929). Pleiotropic loci may be used less frequently by evolution due to negative impacts of changes to one or more of the traits influenced by the pleiotropic gene (Fisher, 1930; Orr, 2000; Otto, 2004; Wright, 1939). Alternatively, pleiotropy could be a driver of evolution if altering a suite of traits results in positive fitness consequences (Mackay and Anholt, 2024; Rennison and Peichel, 2022). Further, pleiotropic genes may be utilized by evolution if the positive phenotypic benefits of altering one trait exceed the negative or neutral consequences of altering other traits (Jeffery, 2005; Wright, 1929). These complexities make it critical to understand the phenotypic consequences of genetic variation at pleiotropic loci, as well as how these loci contribute to the fitness of organisms.

The Mexican cavefish, *Astyanax mexicanus*, has emerged as a leading model to study the genetic basis of trait evolution. A. *mexicanus* is a freshwater species of teleost fish that exists in a riverine surface form and a cave form that inhabits at least 35 caves in Northeastern Mexico (Espinasa et al., 2020; Miranda-Gamboa et al., 2023; Mitchell et al., 1977). The cave form of *A. mexicanus* exhibits drastic morphological differences relative to surface fish, including eye regression and reduced or absent melanin pigmentation (Jeffery, 2001; Jeffery et al., 2003; Şadoğlu, 1957; Sadoglu and McKee, 1969; Wilkens, 1988). The cave forms also exhibit multiple derived behavioral adaptations, including reduced sleep and reductions in social behaviors, enhanced vibration attraction, and altered larval and adult feeding behaviors (Aspiras et al., 2015; Duboué et al., 2011; Elipot et al., 2013; Kowalko et al., 2013a; Lloyd et al., 2018; Patch et al., 2022; Paz et al., 2023; Rodriguez-Morales et al., 2022; Yoshizawa et al., 2010). The diverse number of behavioral and physiological differences between cave and surface fish suggests changes in multiple traits were required for adaptation to the cave environment.

Whether the trait alterations evolved in cavefish are due to the same or distinct genetic loci has been studied extensively (Kowalko et al., 2013a; Kowalko et al., 2013b; O’Gorman et al., 2021; Oliva et al., 2022; Protas et al., 2008; Yamamoto et al., 2009; Yoshizawa et al., 2012). Both manipulation of differentially expressed genes and the presence of overlapping QTL for distinct phenotypes in *A. mexicanus* suggest that pleiotropy may play a role in cavefish evolution (Kowalko et al., 2013a; O’Quin and McGaugh, 2016; Protas et al., 2008; Yamamoto et al., 2009; Yoshizawa et al., 2012). However, few causative loci for cave-evolved traits have been identified in this species, presenting a challenge to distinguishing between pleiotropy and alternative hypotheses, such as closely linked genes contributing to the evolution of different traits in cavefish populations.

One notable exception is albinism. Albinism has evolved repeatedly in cave organisms, and albinism in *A. mexicanus* cavefish is the result of mutations in a single gene, *oculocutaneous albinism II*, or *oca2* (Culver and Pipan, 2019; Klaassen et al., 2018; Protas et al., 2006; Şadoğlu, 1957; Warren et al., 2021). However, in addition to albinism, *oca2* has been implicated in the evolution of other cave-evolved traits, including enhancement of catecholamine levels, anesthesia resistance, and reductions in sleep, suggesting a pleiotropic role for *oca2* in cavefish evolution (Bilandžija et al., 2013; Bilandžija et al., 2018; O’Gorman et al., 2021). While *oca2* alleles are under positive selection in multiple *A. mexicanus* cave populations (O’Gorman et al., 2021), whether *oca2* affects other cave-evolved traits, and which of these trait(s) affected by *oca2* in cavefish provide a benefit in a cave environment are currently unknown.

Here, we investigate the role of mutation of *oca2* in another behavior, prey capture behavior. Previous work has shown that larval surface fish hunt using visual cues under lighted conditions, whereas cavefish capture prey using their lateral line (Lloyd et al., 2018). When hunting in the dark, cave and surface fish use their lateral line to strike prey at wider angles compared to surface fish in lighted conditions (Lloyd et al., 2018). Here, we find that albino, homozygous mutant *oca2* surface fish larvae utilize an altered, cave-like wide strike angle when capturing prey. However, unlike cavefish, *oca2* mutant surface fish utilize this wide-angle striking even when they are using visual cues to capture prey. Finally, we find that albino surface fish show reduced performance in a competition assay against their pigmented siblings under lighted conditions. This pigmented surface fish advantage is absent in dark conditions, raising the possibility that cave alleles of *oca2* provide a disadvantage when foraging under lighted conditions that is absent in dark conditions like those found in caves. Together, this work suggests that *oca2* contributes to the evolution of multiple behavioral traits, some of which may result in decreased fitness in a surface habitat.

## 2. Results

To determine whether loss of OCA2 contributes to the evolution of prey capture behavior, we compared the strike dynamics of gene edited *oca2* mutant surface fish that are homozygous for a two base pair deletion in exon 21 of the *oca2* gene (Klaassen et al., 2018; O’Gorman et al., 2021) to wild-type siblings and Pachón cavefish. Larval fish between 29 and 33 dpf were recorded while eating brine shrimp in a lighted arena and strike attempts were analyzed. Consistent with prior results, the wild-type surface fish larvae strike prey head on, frequently exhibiting a J turn movement, while Pachón cavefish approach prey from the side, frequently using a C turn movement to capture prey (Lloyd et al., 2018) (Fig. 1A). Unlike their wild-type siblings, the albino, *oca2* mutant (*oca2*^*Δ2bp/Δ2bp*^) surface fish larvae exhibited a large strike angle when feeding, similar to what is observed in cavefish (Fig. 1A). Quantification of the distance between the fish and the prey prior to striking revealed no significant differences amongst any populations (Fig. 1B). However, the total number of attempted strikes and the proportion of successful strikes were both reduced in *oca2*^*Δ2bp/Δ2bp*^ surface larvae compared to wild-type surface larvae (Fig. 1C&D). Together, these results demonstrate that *oca2* mutant surface fish display shifts in multiple components of larval prey capture compared to wild-type surface fish in the light, and they display prey capture behavior similar to Pachón cavefish.

**Figure 1.**
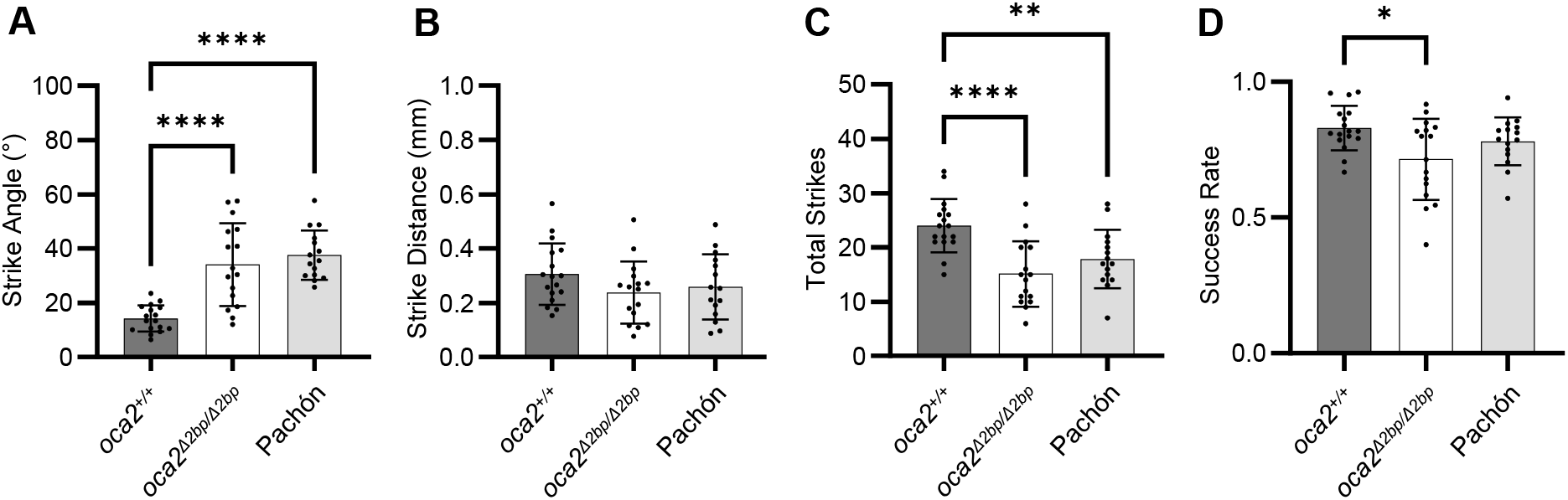
oca2 mutant surface fish exhibit altered strike dynamics in the light. (A) Average strike angles of wild-type (*oca2*^*+/+*^) surface fish (n = 17), *oca2*^*Δ2bp/Δ2bp*^ surface fish (n =16), and Pachón cavefish (n = 15). One way ANOVA, F(2, 45) = 23.30, p <0.0001. Tukey’s multiple comparisons test: *oca2*^*+/+*^ vs *oca2*^*Δ2bp/Δ2bp*^ surface fish (Adjusted p <0.0001), *oca2*^*+/+*^ surface vs Pachón (Adjusted p <0.0001), *oca2*^*Δ2bp/Δ2bp*^ vs Pachón (Adjusted p = 0.6398). (B) Average strike distances of *oca2*^*+/+*^ surface fish (n = 17), *oca2*^*Δ2bp/Δ2bp*^ surface fish (n = 16), and Pachón cavefish (n = 15). One way ANOVA, F(2, 45) = 1.521, p = 0.2296. (C) Total number of strikes for *oca2*^*+/+*^ surface fish (n = 17), *oca2*^*Δ2bp/Δ2bp*^ surface fish (n = 16), and Pachón cavefish (n = 15). One way ANOVA, F(2, 45) = 11.54, p <0.0001. Tukey’s multiple comparisons test: *oca2*^*+/+*^ vs *oca2*^*Δ2bp/Δ2bp*^ surface fish (Adjusted p<0.0001), *oca2*^*+/+*^ surface fish vs Pachón (Adjusted p = 0.0074), *oca2*^*Δ2bp/Δ2bp*^ vs Pachón (Adjusted p = 0.3488). (D) Success rates of wild-type surface fish (n = 17), *oca2*^*Δ2bp/Δ2bp*^ surface fish (n = 16), and Pachón cavefish (n = 15). One way ANOVA, F(2, 45) = 4.445, p = 0.0173. Tukey’s multiple comparisons test: *oca2*^*+/+*^ vs *oca2*^*Δ2bp/Δ2bp*^ surface fish (Adjusted p = 0.0128), *oca2*^*+/+*^ surface fish vs Pachón (Adjusted p = 0.4285), *oca2*^*Δ2bp/Δ2bp*^ vs Pachón (Adjusted p = 0.2363). All error bars are standard error of the mean. ^*^p<0.05, ^**^p<0.01, ^***^p<0.001, ^****^p<0.0001

As surface fish alter their prey capture behavior in the dark (Lloyd et al., 2018), we next performed prey capture assays in both light and dark conditions to determine if *oca2* mutant surface fish also alter their behavior in the dark. Wild-type surface fish exhibit an increase in strike angle and no change in strike distance in dark conditions when compared to light conditions (Fig. 2A-B, S1A-B). Further, wild-type surface fish attempted to strike less frequently, and showed a decrease in success rate under dark conditions (Fig. 2C-D, S1C-D). In contrast, *oca2*^*Δ2bp/Δ2bp*^ surface fish demonstrated no change in any metric between light and dark conditions (Fig. 2A-D), similar to Pachón cavefish (Figure S1A-D). Together, these data suggest that *oca2*-mutant surface fish do not alter their feeding dynamics in the absence of visual cues and raise the possibility that *oca2*^*Δ2bp/Δ2bp*^ surface larvae may be using vision-independent methods of feeding under lighted conditions.

**Figure 2.**
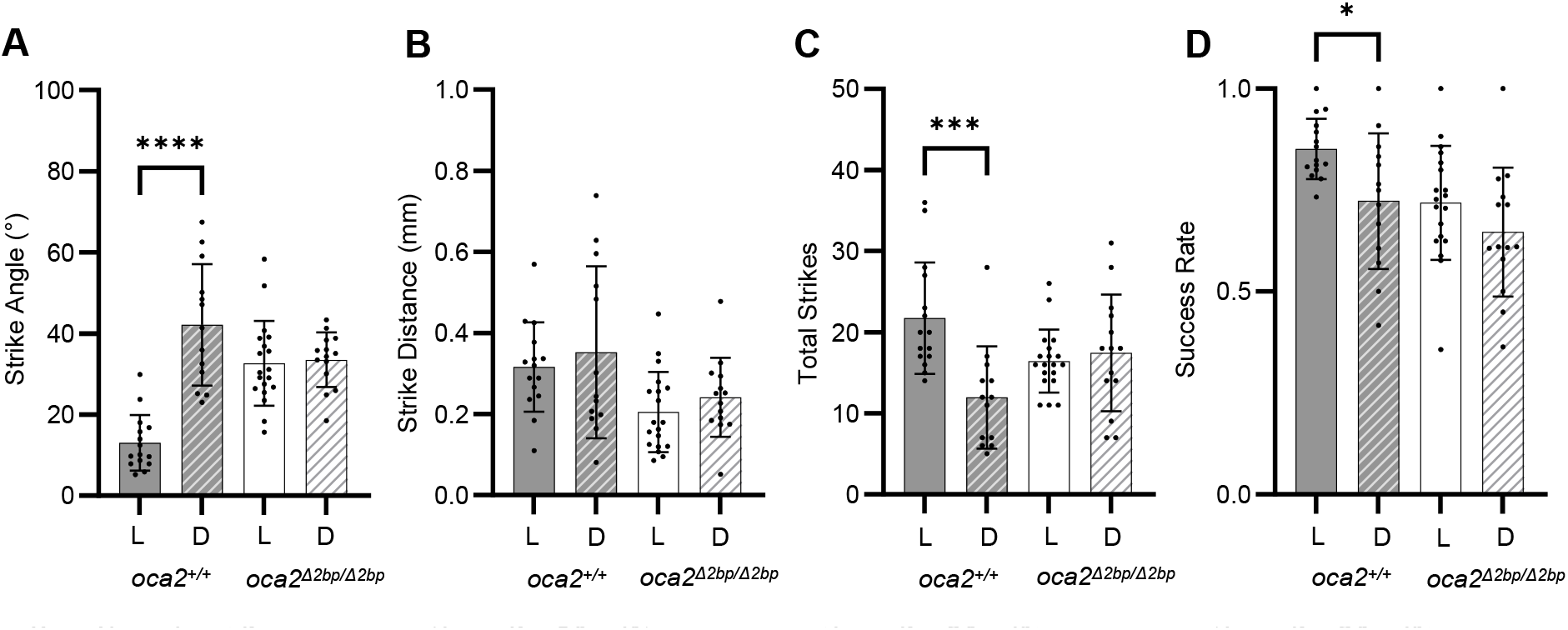
oca2 mutant surface fish do not alter their strike dynamics in the dark. (A) Average strike angles of wild-type (*oca2*^*+/+*^) *oca2*^*Δ2bp/Δ2bp*^ surface fish in light and dark conditions. Welch’s t test: oca2^+/+^, light, (n = 15) and dark (n = 13), p<0.0001. t test: *oca2*^*Δ2bp/Δ2bp*^, light (n = 19) and dark (n = 14), p = 0.7862. (B) Average strike distances of wild-type and *oca2*^*Δ2bp/Δ2bp*^ surface fish in light and dark conditions. Welch’s t test: oca2^+/+^, light, (n = 15) and dark (n = 13), p = 0.5856. t test: *oca2*^*Δ2bp/Δ2bp*^, light (n = 19) and dark (n = 14), p = 0.3007. (C) Total number of strikes in wild-type and *oca2*^*Δ2bp/Δ2bp*^ surface fish in light and dark conditions. Mann-Whitney U test: oca2^+/+^, light, (n = 15) and dark (n = 13), p = 0.0001. Welch’s t test: *oca2*^*Δ2bp/Δ2bp*^, light (n = 19) and dark (n = 14), p = 0.6401. (D) Success rates of wild-type and *oca2*^*Δ2bp/Δ2bp*^ surface fish in light and dark conditions. Welch’s t test: *oca2*^*+/+*^, light, (n = 15) and dark (n = 13), p = 0.0208. t test: *oca2*^*Δ2bp/Δ2bp*^, light (n = 19) and dark (n = 14), p = 0.1804. All error bars are standard error of the mean. ^*^p<0.05, ^**^p<0.01, ^***^p<0.001, ^****^p<0.0001

One possibility for the striking behaviors observed in *oca2* mutant surface fish is that these fish are incapable of feeding using visual cues due to visual system defects. We quantified optomotor response, an innate behavior characterized by fish swimming in the same direction as a moving visual stimulus, which has been used previously in zebrafish to assess for loss of visual function (Clark, 1981; Neuhauss, 2003; Neuhauss et al., 1999). Fish were placed in a rectangular well, exposed to high contrast moving lines, and scored for the proportion of the distance traveled across the well in the direction of the moving lines following each directional switch. Fish with an optomotor response index (OMI) close to 1 frequently traveled in the direction of the lines whereas fish with an OMI close to zero moved without regard to the movement of the lines. Surface fish show a robust optomotor response (Fig. S2A). In contrast, Pachón cavefish do not swim in the direction of the moving lines and have OMI around zero (Fig. S2A), suggesting they do not display an optomotor response and do not have visual function. Similar to wild-type surface fish siblings, the majority of *oca2*^*Δ2bp/Δ2bp*^ surface fish have OMI close to 1, indicating that they respond robustly in this assay, and suggesting they retain visual function (Fig. S2B).

The presence of robust visually guided behavior in *oca2* mutant surface fish raises the possibility that these fish are performing wide strike angles while feeding using visual cues. Alternatively, *oca2* mutant surface fish may preferentially feed using lateral line cues. In order to test what sensory modality *oca2* mutant fish use while hunting, we ablated the lateral line using gentamicin, an ototoxic compound used in fish species (Song et al., 1995; Van Trump et al., 2010), and assayed prey capture under light and dark conditions. Wild-type surface larvae, in lighted conditions, demonstrated no significant difference in strike angle, distance, total strikes, or success rate between gentamicin treated and untreated groups, and feed at low strike angles associated with visual-based feeding (Fig 3A-D). In contrast, in lighted conditions, *oca2*^*Δ2bp/Δ2bp*^ surface fish strike at wide angles regardless of gentamicin treatment. Further, strike angle increases when the lateral line is ablated following gentamicin treatment in *oca2*^*Δ2bp/Δ2bp*^ surface fish (Fig 3A). This suggests that *oca2* mutant surface larvae do not require lateral line-mediated cues for wide-angle strikes in lighted conditions. While *oca2*^*Δ2bp/Δ2bp*^ surface fish showed no significant differences in strike distance and success rate between treated and untreated groups in lighted conditions, they did show a decrease in total strike attempts (Fig 3B-D), suggesting that the lateral line may play a role in finding prey in these fish under lighted conditions. In the dark, neither wild-type or *oca2*^*Δ2bp/Δ2bp*^ surface fish were able to capture prey when gentamicin-treated, indicating that *oca2* mutant surface fish require either vision or the lateral line for hunting, similar to their wild-type counterparts (Fig. 3). Together, these results suggest that the wide angle used during hunting by *oca2* mutant surface fish is not due to a shift to lateral line mediated feeding under lighted conditions, but instead occurs in surface fish using vision to hunt.

**Figure 3.**
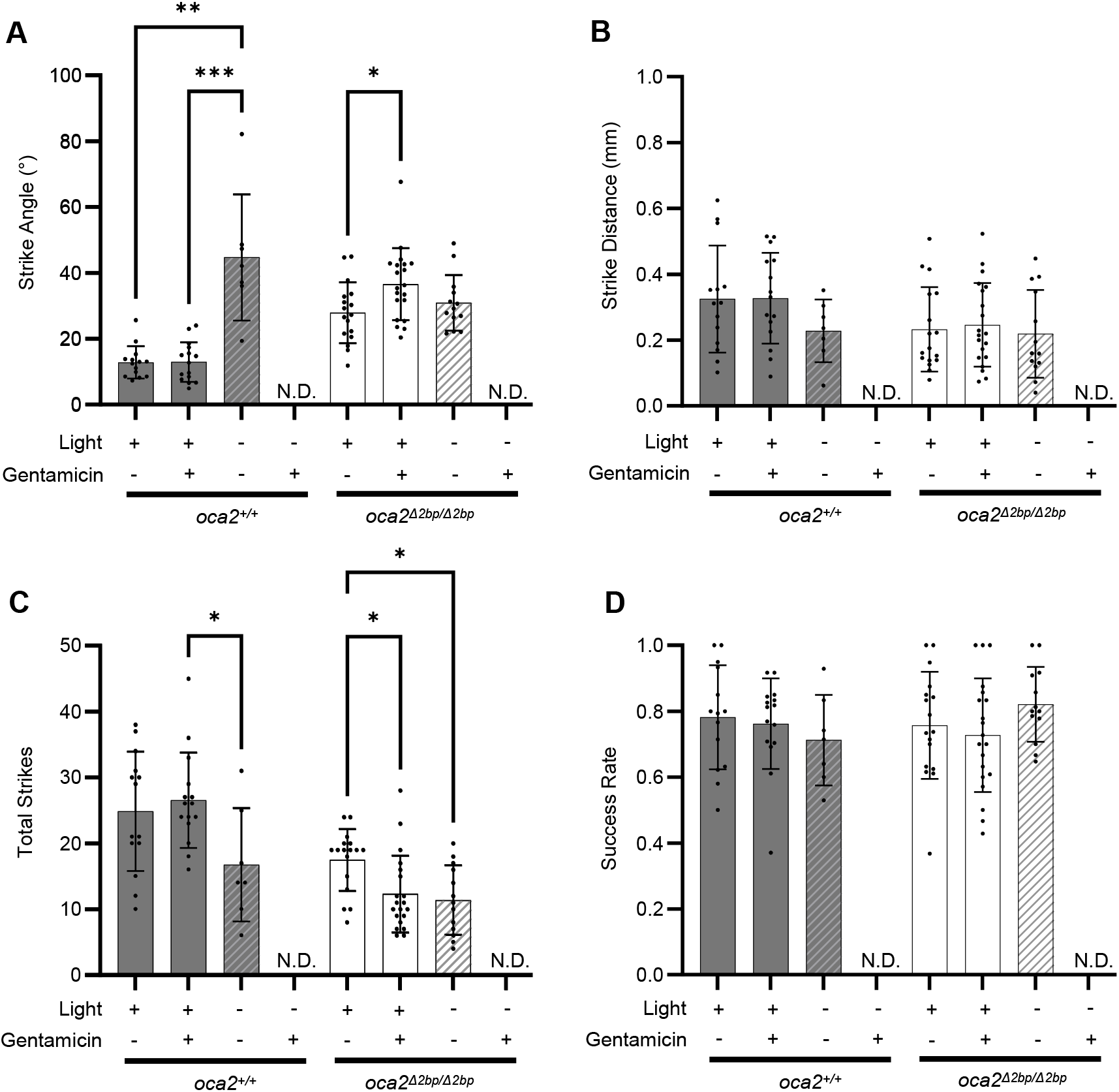
oca2 mutant fish strike at wide angles after lateral line ablation. (A) Strike angles of wild type (*oca2*^*+/+*^) and *oca2*^*Δ2bp/Δ2bp*^ surface fish, in light and dark conditions and with or without gentamicin treatment. *oca2*^*+/+*^ surface fish, light, untreated (n = 14), *oca2*^*+/+*^ surface fish, light, treated (n = 15), *oca2*^*+/+*^ surface fish, dark, untreated (n = 7), and *oca2*^*+/+*^ surface fish, dark, treated (No Data). Kruskal-Wallis test, number of treatments = 3, number of values = 36, KW statistic = 15.51, p = 0.0004. Dunn’s multiple comparisons test: light/untreated vs light/treated (Adjusted p >0.9999), light/untreated vs dark/untreated (Adjusted P = 0.0012), light/treated vs dark/untreated (Adjusted p = 0.0008). *oca2*^*Δ2bp/Δ2bp*^ surface fish, light, untreated (n = 17), *oca2*^*Δ2bp/Δ2bp*^ surface fish, light, treated (n = 20), *oca2*^*Δ2bp/Δ2bp*^ surface fish, dark, untreated (n =13), and *oca2*^*Δ2bp/Δ2bp*^ surface fish, dark, treated (No Data). One way ANOVA, F (2, 47) = 3.765, p = 0.0304. Tukey’s multiple comparisons test: light/untreated vs light/treated (Adjusted p = 0.0261), light/untreated vs dark/untreated (Adjusted p = 0.6815), light/treated vs dark/untreated (Adjusted p = 0.2449). (B) Average strike distances of wild-type and *oca2*^*Δ2bp/Δ2bp*^ surface fish, in light and dark conditions and with or without gentamicin treatment. *oca2*^*+/+*^ surface fish, light, untreated (n = 14), *oca2*^*+/+*^ surface fish, light, treated (n = 15), *oca2*^*+/+*^ surface fish, dark, untreated (n = 7), and *oca2*^*+/+*^ surface fish, dark, treated (No Data). One way ANOVA, F (2, 33) = 1.341, p = 0.2756. *oca2*^*Δ2bp/Δ2bp*^ surface fish, light, untreated (n = 17), *oca2*^*Δ2bp/Δ2bp*^ surface fish, light, treated (n = 20), *oca2*^*Δ2bp/Δ2bp*^ surface fish, dark, untreated (n = 13), and *oca2*^*Δ2bp/Δ2bp*^ surface fish, dark, treated (No Data). Kruskal-Wallis test, number of treatments = 3, number of values = 50, KW statistic = 0.5835, p = 0.7469. (C) Total number of strikes for wild-type and *oca2*^*Δ2bp/Δ2bp*^ surface fish, in light and dark conditions and with or without gentamicin treatment. *oca2*^*+/+*^ surface fish, light, untreated (n = 14), *oca2*^*+/+*^ surface fish, light, treated (n = 15), *oca2*^*+/+*^ surface fish, dark, untreated (n = 7), and *oca2*^*+/+*^ surface fish, dark, treated (No Data). One way ANOVA, F (2, 33) = 3.495, p = 0.0420. Tukey’s multiple comparisons test: light/untreated vs light/treated (Adjusted p = 0.8495), light/untreated vs dark/untreated (Adjusted p = 0.0998), light/treated vs dark/untreated (Adjusted p = 0.0365). *oca2*^*Δ2bp/Δ2bp*^ surface fish, light, untreated (n = 17), *oca2*^*Δ2bp/Δ2bp*^ surface fish, light, treated (n = 20), *oca2*^*Δ2bp/Δ2bp*^ surface fish, dark, untreated (n = 13), and *oca2*^*Δ2bp/Δ2bp*^ surface fish, dark, treated (No Data). Kruskal-Wallis test, number of treatments = 3, number of values = 50, KW statistic = 11.38, p = 0.0034. Dunn’s multiple comparisons test: light/untreated vs light/treated (Adjusted p = 0.0111), light/untreated vs dark/untreated (Adjusted p = 0.0109), light/treated vs dark/untreated (Adjusted p >0.9999). (D) Success rates of wild type and *oca2*^*Δ2bp/Δ2bp*^ surface fish, in light and dark conditions and with or without gentamicin treatment. *oca2*^*+/+*^ surface fish, light, untreated (n = 14), *oca2*^*+/+*^ surface fish, light, treated (n = 15), *oca2*^*+/+*^ surface fish, dark, untreated (n = 7), and *oca2*^*+/+*^ surface fish, dark, treated (No Data). Kruskal-Wallis test, number of treatments = 3, number of values = 36, KW statistic = 1.171, p = 0.5569. *oca2*^*Δ2bp/Δ2bp*^ surface fish, light, untreated (n = 17), *oca2*^*Δ2bp/Δ2bp*^ surface fish, light, treated (n = 20), *oca2*^*Δ2bp/Δ2bp*^ surface fish, dark, untreated (n = 13), and *oca2*^*Δ2bp/Δ2bp*^ surface fish, dark, treated (No Data). One way ANOVA, F (2, 47) = 1.437, P = 0.2478. All error bars are standard error of the mean. ^*^p<0.05, ^**^p<0.01, ^***^p<0.001, ^****^p<0.0001

We next sought to determine whether these differences in hunting behavior provided an advantage to fish under conditions that mimic surface and cave environments. To determine whether surface or cave-like *oca2* alleles provide an advantage while hunting, we developed a competition assay, in which two fish were provided with a small number of prey, and we determined what proportion of the prey was eaten by individuals of different genotypes (Fig 4A). We found that, on average, surface fish consume more prey than cavefish when competing under lighted conditions, and cavefish consumed more prey than surface fish in the dark (Fig 4B). While we found that pigmented fish on average outcompeted albino siblings under lighted conditions, we found no significant differences between albino and pigmented siblings under dark conditions (Fig 4C). Together, these results suggest that mutations in *oca2* result in a disadvantage for surface larvae when hunting in the light, and that this disadvantage is alleviated under dark conditions.

**Figure 4.**
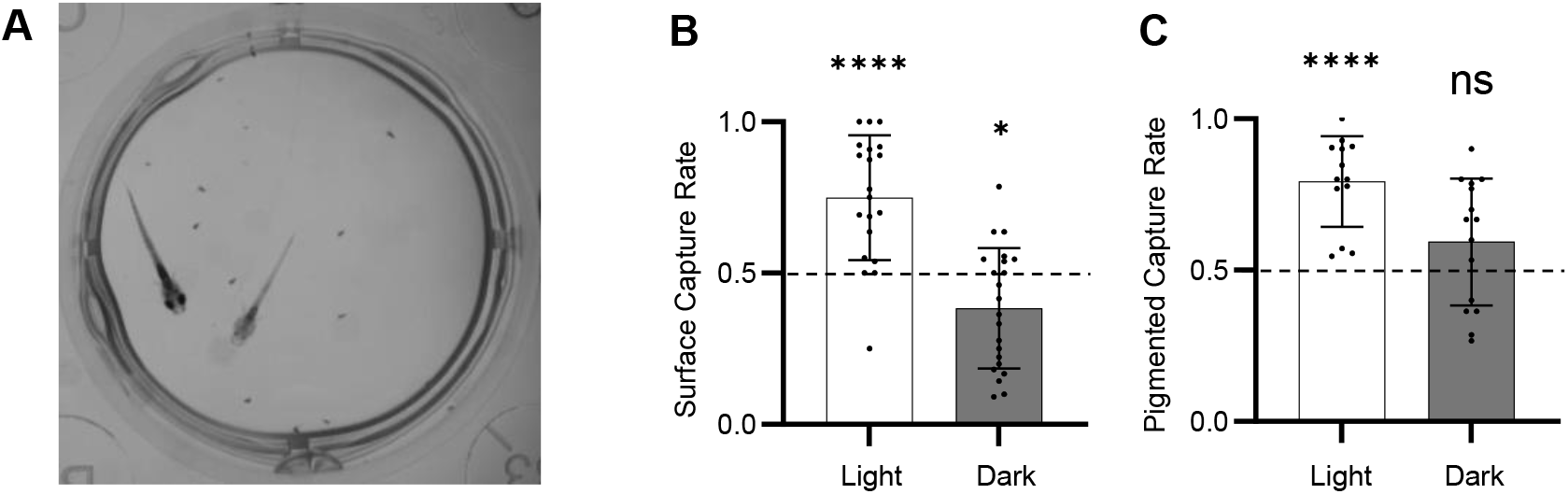
Competition assays reveal differences in hunting success between populations. (A) Representative image of competition assay with a surface fish and Pachón cavefish. (B) Proportion of total prey captured by surface fish competitors in a competition assay between a surface fish and Pachón cavefish in light and dark conditions. One-Sampled T test, n = 20 trials (light), 22 (dark), theoretical mean = 0.5, actual means = 0.7492 (light), 0.3841 (dark), t = 5.405 (light), 2.728 (dark), df = 19 (light), 21 (dark), p <0.0001 (light), 0.0126 (dark). (C) Proportion of totally prey captured by pigmented surface fish sibling competitors (*oca2*^*Δ2bp/+*^ or *oca*^*+/+*^) in a competition assay between a pigmented surface fish and an *oca2*-mutant albino surface fish in light and dark conditions. One-Sampled T test, n = 13 trials (light), 15 (dark), theoretical mean = 0.5, actual means = 0.7929 (light), 0.5934 (dark), t = 7.053 (light), 1.723 (dark), df = 12 (light), 14 (dark), p = <0.0001 (light), 0.1069 (dark). All error bars are standard error of the mean. ^*^p<0.05, ^**^p<0.01, ^***^p<0.001, ^****^p<0.0001

## 3. Discussion

Here, we quantified larval feeding behavior in surface fish with mutations in the *oca2* gene, a pleiotropic gene that is responsible for albinism and contributes to sleep loss in multiple *A. mexicanus* cavefish populations (Klaassen et al., 2018; O’Gorman et al., 2021; Protas et al., 2006) to determine if *oca2* has other pleiotropic effects in this species. We found that *oca2*^*Δ2bp/Δ2bp*^ larvae exhibit altered prey capture behavior compared to wild-type, pigmented siblings, and feed at wider strike angles. This altered behavior is not due to complete loss of visual function in *oca2* mutant larvae and a subsequent shift to lateral line-mediated feeding, as these fish have an optomotor response similar to wild-type siblings. Further supporting that *oca2* mutant surface fish hunt with wide strike angles using visual cues, when we ablated the lateral line of *oca2* mutant surface fish, these fish continued to feed with wide string angles. Finally, we assessed whether loss-of-function *oca2* alleles provide a benefit to feeding in a habitat that is dark, similar to the cave. We found that while pigmented siblings outcompeted *oca2* mutant larvae in light conditions, there was no significant difference in successful hunting under dark conditions, suggesting that *oca2* is important for larval feeding behavior under light, but not dark conditions.

### Pleiotropy: Pigmentation and Behavior

Genes that impact pigmentation have been associated with differences in behavior across taxa (Ducrest et al., 2008; Reissmann and Ludwig, 2013). For example, genetic variation at the agouti locus impacts both pigmentation and aggressive behavior in mice (Carola et al., 2014). Fruit flies with a mutation in the *ebony* gene exhibit higher aggression and increased sleep compared to wild-type files (Pantalia et al., 2023). Additionally, albinistic animals within a species can exhibit different behaviors than their pigmented counterparts. Albino rats have altered sleep and aggression, and albino catfish (*Siluris glanis*) exhibited lower aggression and reduced shoaling behavior compared to pigmented catfish (Barnett and Hocking, 1981; Barnett et al., 1979; Slavík et al., 2016). Together, these studies suggest that pleiotropic loci can impact both pigmentation and behavior.

In *A. mexicanus*, pleiotropy at the *oca2* locus has been proposed to play a role in the evolution of both pigmentation and behavior (Bilandžija et al., 2013; Bilandžija et al., 2018; O’Gorman et al., 2021). Coding mutations at the *oca2* locus are responsible for albinism in at least two *A. mexicanus* cavefish populations (Klaassen et al., 2018; Protas et al., 2006). Further, mutations in *oca2* have been proposed to contribute to increases in catecholamine levels found in cavefish, as well as increases in anesthesia resistance and reductions in sleep (Bilandžija et al., 2013; Bilandžija et al., 2018; O’Gorman et al., 2021). Here, we identify another potential role for *oca2* in the evolution cavefish behavior: alterations to larval prey capture behaviors. Together, these studies strongly suggest a role for pleiotropy in the evolution of pigmentation and behavior in cavefish. While albinism is controlled by a single gene in cavefish, reductions in melanin pigmentation in cavefish populations are multigenic (Protas et al., 2006; Protas et al., 2007; Şadoğlu, 1957; Sadoglu and McKee, 1969; Wilkens, 1988). Whether natural variation at other loci in *A. mexicanus* contributes to both pigmentation and behavior is currently unknown.

### The effects of albinism on visual acuity

*A. mexicanus* surface larvae primarily use visual cues for hunting (Lloyd et al., 2018), thus it is important to confirm that the alterations to *oca2* mutant surface larvae feeding behavior are not due to poor visual acuity. Albinism is known to confer visual defects across taxa. Oculocutaneous albinism in humans is highly comorbid with strabismus, nystagmus, foveal hypoplasia, and reduced visual acuity (Grønskov et al., 2007). Albino model organisms also exhibit vision defects. Albino rats, albino mice, and hypopigmented zebrafish have reduced visual acuity when compared to pigmented counterparts (Braha et al., 2021; Li et al., 2022; Ren et al., 2002). Together, these data suggest that albinism’s impact on vision is highly conserved. Our optomotor response data demonstrated that *oca2* mutant larvae are not blind (Fig. S2B) and ablation of the lateral line in these fish suggests that even when hunting using only visual cues, the *oca2* mutant larvae still hunt at wide angles, unlike wild-type siblings (Fig. 3A). Together, these results suggest that *oca2* mutant surface fish strike at prey from wide angles even when using vision. However, we cannot rule out that *oca2* mutant larvae have more subtle visual defects which impact hunting behavior in these animals.

### The adaptive value of oca2: A role for feeding?

The *oca2* gene has previously been implicated in multiple other traits that have evolved in cavefish, including albinism, sleep, and anesthesia resistance (Bilandžija et al., 2018; Klaassen et al., 2018; O’Gorman et al., 2021; Protas et al., 2006). Further, the *oca2* locus is under positive selection in surface fish and multiple cavefish populations (O’Gorman et al., 2021). While there are evolutionary benefits to having pigmentation in a lighted environment, such as camouflage and UV protection (Lin and Fisher, 2007; Sköld et al., 2016), the benefit of loss of OCA2 in the cave habitat is currently unknown. Here, we tested whether having *oca2* mutant alleles provides a benefit to surface fish when hunting in a dark environment. We found Pachón larvae outperformed surface larvae in dark, and surface larvae outcompeted Pachón larvae in the light, suggesting that fish from each population have adapted to hunting in their respective environment conditions, similar to previous studies performed in larval and adult fish (Espinasa et al., 2014; Yoshizawa et al., 2010). However, the *oca2* mutant larvae did not exhibit a significant difference in competition with their pigmented siblings in dark conditions. In contrast, pigmented siblings significantly outperformed their albino siblings under lighted conditions, indicating that albinism negatively effects hunting behavior in the light. These data suggest *oca2*-mediated changes in hunting behavior likely do not provide a competitive benefit in the dark. However, they reveal a potential benefit for functional OCA2 during hunting in the light.

These data raise the possibility that the effects of loss of *oca2* on hunting behavior does not have a negative or positive effect on fish living in the complete darkness of the cave environment. Instead, a dark environment may remove the fitness advantage provided by pigmentation via an intact OCA2 when hunting in the light. This could allow *oca2* to incur mutations without an immediate negative fitness change on this behavior, thus enabling *oca2* to be selected for its other pleiotropic effects in cavefish.

## 4. Materials and Methods

### 4.1 Fish Husbandry and Populations

Fish were bred and larvae were raised as previously described (Borowsky, et al., 2008b, Kozol, et al., 2022). All fish were housed under a 14:10 light/dark cycle. Larvae were kept at 25°C for the first 6 days post fertilization (dpf) in glass bowls, and then transferred to tanks, where larvae were raised at a density of 30 fish per 2 liter tank, with water changes twice per week at 23°C. Larvae were fed *Artemia salina* to satiation twice per day, starting at 6 dpf. Larvae were not fed in the afternoon before prey capture assays or competition assays, to ensure satiety was not met before experiments.

Surface fish lines with a mutant *oca2* allele were derived previously, and fish assayed here had a 2 bp deletion in exon 21, the exon that is absent in Molino cavefish populations, of the *oca2* gene (Klaassen et al., 2018; O’Gorman et al., 2021). Surface fish heterozygous at the *oca2* locus were incrossed to produce albino (*oca2* mutant) and pigmented (heterozygous and homozygous wild-type, referred to as wild-type) offspring. Albino, *oca2* mutant fish were compared to wild-type siblings for all strike assays, and to pigmented siblings (wild-type or heterozygous) for competition assays. Wild-type fish were distinguished from heterozygous siblings by genotyping following assays. Surface and cavefish populations used in this study were derived from lab-bred populations captured multiple generations ago in Mexico, and in the case of cavefish, from the Pachon cave.

### 4.2 Prey Capture Assays

All larvae for prey capture assays were between 29-33 dpf. One well in an untreated 12-well plate (24.0mm in diameter) served as the arena. 25-30 *Artemia* were added to the arena immediately before the assay. All *Artemia* used were prepared 24 hours prior to experimentation, so that all prey were of similar sizes and stages of development across trials. *Artemia* were prepared by placing cysts in saltwater (35g/L) with constant aeration. Hatched *Artemia* were separated from cysts and transferred to fresh fish system water (pH: 7.0-7.3, conductivity: 600-800μS/cm) prior to assays.

Larvae were placed in the arena and feeding was assessed for 2 minutes following addition of the larva. Videos were recorded at 50 frames per second (fps) and a resolution of 992×1000 using an FLIR Grasshopper3 High Performance USB 3.0 Monochrome Camera (Edmund Optics Cat. No. 88-607) with a 12mm HP series lens, 1/1” (Edmund Optics, Cat. No. 86570). Videos were recorded using the program Spinview, from FLIR’s Spinnaker SDK. Prey capture assays in lighted conditions were backlit via white LED strip lights, and filmed from above. Overhead lights were turned off during experimentation to prevent glare on the camera. For assays in dark conditions, the arena was placed in an opaque box illuminated with IR lights with a blackout curtain draped over top. The arena was filmed from below to increase contrast of prey and fish.

All larval strikes within 2 minutes of the larvae entering the arena were recorded. Strikes were broken into three categories: unsuccessful strikes, successful strikes, unmeasurable successful strikes. Unsuccessful strikes were strike attempts from the larvae that did not result in successfully capturing *Artemia*. Successful strikes were strike attempts where the larvae captured *Artemia*. Unmeasurable successful strikes were strikes where the larva captured prey, but the strike could not be quantified for angle or distance, which were usually the result of the larvae being rotated on its side, or when the larvae performed a “multi-strike.” A multi-strike is where the larvae performed multiple strike attempts in quick succession with no recovery between strikes. Angle and distance could not be quantified as the starting position was not reflective of the capturing position. Unmeasurable strikes were omitted from angle or distance measurements, but still counted towards total strikes or success rate.

Successful strikes were quantified for strike angle and strike distance by measuring in FIJI (Schindelin et al., 2012), as previously described (Lloyd et al., 2018). Briefly, on the frame before the initiation of the strike, the shortest distance between the *Artemia* and the larvae’s head was measured. Additionally, the angle between the midline of the larvae from the base of its head and the center of the prey was measured to obtain the strike angle. Prey capture was further analyzed for total of number of strikes and success rate. Unmeasurable strikes, such as larvae-rotated strikes or multi-strikes, were included in the quantification of success rate.

### 4.3. Optomotor Assays

All larvae for OMR assays were 8 dpf. Larvae were placed in a 4-well rectangular arena (Nunc rectangular dishes, Thomas Scientific, item number 1228D90). A Samsung Tab Active Pro tablet (model number SM-T540) was positioned below the arena, providing backlight and a video playing the moving lines. The video consists of a 30 second white background followed by alternating 2 centimeter black & white lines moving at a speed of 1 centimeter per second. The moving lines switched direction 5 times every 30 seconds, ending on another 30 second white screen. OMR Assays were filmed from above using an FLIR Grasshopper3 High Performance USB 3.0 Monochrome Camera (Edmund Optics Cat. No. 88-607) with a 12mm HP series lens, 1/1” (Edmund Optics, Cat. No. 86570). Videos were recorded using the program Spinview, from FLIR’s Spinnaker SDK. Recording resolution was 800×1200 and framerate was 30 fps.

OMR videos were analyzed via FIJI (Schindelin et al., 2012). Scales were set by using the line tool drawn to the total length of the well, 78mm. The X-position of each larva was taken during the line switch; if larvae were obscured at that point, their most recent known position was used instead. The recorded positions were subtracted from one another to assess what distance each larva swam in the 30 second interval in the direction of the moving lines. The first interval was not recorded as the larvae’s starting position had not been influenced by moving lines. All other intervals were averaged together and converted to a percentage to gauge the performance of each larva’s optomotor response, recorded as the optomotor index.

### 4.4. Competition Assays

All larvae for competition assays were 29-33 dpf. Surface fish and Pachon cavefish were age-matched. Pigmented *oca2* fish (*oca2*^*Δ2bp/+*^ and *oca*^*+/+*^) competed against their albino (*oca2*^*Δ2bp/Δ2bp*^) siblings. After a two-minute acclimation, 8-14 *Artemia* were added to the arena. Recording continued until 0-1 *Artemia* remained. Videos were recorded from above on an FLIR Grasshopper3 High Performance USB 3.0 Monochrome Camera (Edmund Optics Cat. No. 88-607) with a 50mm HP series lens, 4/3” (Edmund Optics, Cat. No. 86574) at 50fps and a resolution of 992×1000.

Competition assay recordings were analyzed for total number of *Artemia* added and how many *Artemia* each larvae captured. Recordings where less than 8 or more than 14 *Artemia* were added were omitted to ensure similar levels of total prey for comparisons. Total *Artemia* captured were converted into a percentage per focal fish (wild-type surface or pigmented) based off the starting amount of *Artemia*.

### 4.5. Gentamicin treatment

Larvae had their lateral line ablated via treatment of 0.001% gentamicin sulfate solution 24 hours before prey capture assays in line with previous methods established in *A. mexicanus* (Lloyd et al., 2018). Lateral line ablation was validated on a subset of fish from each trial via visualization of neuromasts following 20μg/ml DASPEI solution staining for 20-25 minutes. DASPEI staining was performed in darkness, to prevent degradation of fluorescence.

### 4.6. Genotyping

Lines of surface fish harboring a 2 base pair (bp) mutation at the *oca2* locus (indicated by Δ2bp) were established previously (Klaassen et al., 2018; O’Gorman et al., 2021). Fish heterozygous for the allele were identified by fin clipping, and incrossed to produce pigmented (heterozygous or wild-type) and homozygous mutant offspring. Sibling fish were compared for all assays. The *oca2*^Δ2bp/Δ2bp^ larvae were identified by the lack of pigmentation. Pigmented siblings were genotyped after assaying to determine whether they were *oca2*^+/+^ or *oca2*^Δ2bp/+^ using previously described methods (O’Gorman et al., 2021). Briefly, pigmented larvae were euthanized, then placed in 100μl 50mM NaOH and heated to 95°C for 30 minutes. After heating, 10μl 1M Tris-HCl pH 8.0 was added. Two PCRs were performed per sample using allele-specific forward primers; 5’-CTGGTCATGTGGGTCTCAGC-3’ to test for wild type *oca2*, 5’-TCTGGTCATGTGGGTCTCATT-3’ to test for mutant *oca2*, and the reverse primer for both reactions, 5’-TTTCCAAAGATCACATATCTTGAC-3’ (O’Gorman et al., 2021).

### 4.10. Statistical Analysis

All results were first tested for normal distribution using a Shapiro-Wilkes normality test. Results with non-normal distribution were then compared using a Mann-Whitney U test. Results with normal distribution were then tested for equal variance across groups via Levene’s test. If groups did not have equal variance, then a Welch’s t-test was performed. If equal variances were present, then a student’s t-test was used. Additionally, if more than two samples were compared, results with normal distribution were tested via one-way ANOVA followed by post-hoc test or a Kruskal-Wallis test if not normally distributed. All statistical tests were performed using and graphs generated with Graphpad Prism version 10.0.0, Graphpad Software, graphpad.com.

## Acknowledgements

This work was funded by NSF grant award 2147597 to JEK and ACK and NIH award R35GM138345 to JEK.

## References

Aspiras, A. C., Rohner, N., Martineau, B., Borowsky, R. L. and Tabin, C. J. (2015). Melanocortin 4 receptor mutations contribute to the adaptation of cavefish to nutrient-poor conditions. Proc. Natl. Acad. Sci. U. S. A. 112, 9668–9673.

Barnett, S. A. and Hocking, W. E. (1981). Further experiments on the social interactions of domestic “Norway” rats. Aggress. Behav. 7, 259–263.

Barnett, S. A., Dickson, R. G. and Hocking, W. E. (1979). Genotype and environment in the social interactions of wild and domestic “Norway” rats. Aggress. Behav. 5, 105–119.

Bilandžija, H., Ma, L., Parkhurst, A. and Jeffery, W. R. (2013). A potential benefit of albinism in Astyanax cavefish: downregulation of the oca2 gene increases tyrosine and catecholamine levels as an alternative to melanin synthesis. PLoS One 8, e80823.

Bilandžija, H., Abraham, L., Ma, L., Renner, K. J. and Jeffery, W. R. (2018). Behavioural changes controlled by catecholaminergic systems explain recurrent loss of pigmentation in cavefish. Proc. Biol. Sci. 285,.

Bolnick, D. I., Barrett, R. D. H., Oke, K. B., Rennison, D. J. and Stuart, Y. E. (2018). (Non)Parallel Evolution. Annu. Rev. Ecol. Evol. Syst. 49, 303–330.

Braha, M., Porciatti, V. and Chou, T.-H. (2021). Retinal and cortical visual acuity in a common inbred albino mouse. PLoS One 16, e0242394.

Carola, V., Perlas, E., Zonfrillo, F., Soini, H. A., Novotny, M. V. and Gross, C. T. (2014). Modulation of social behavior by the agouti pigmentation gene. Front. Behav. Neurosci. 8, 259.

Clark, D. T. (1981). VISUAL RESPONSES IN DEVELOPING ZEBRAFISH (BRACHYDANIO RERIO).

Conte, G. L., Arnegard, M. E., Peichel, C. L. and Schluter, D. (2012). The probability of genetic parallelism and convergence in natural populations. Proc. Biol. Sci. 279, 5039– 5047.

Culver, D. C. and Pipan, T. (2019). The Biology of Caves and Other Subterranean Habitats.

Duboué, E. R., Keene, A. C. and Borowsky, R. L. (2011). Evolutionary convergence on sleep loss in cavefish populations. Curr. Biol. 21, 671–676.

Ducrest, A.-L., Keller, L. and Roulin, A. (2008). Pleiotropy in the melanocortin system, coloration and behavioural syndromes. Trends Ecol. Evol. 23, 502–510.

Elipot, Y., Hinaux, H., Callebert, J. and Rétaux, S. (2013). Evolutionary shift from fighting to foraging in blind cavefish through changes in the serotonin network. Curr. Biol. 23, 1–10.

Espinasa, L., Bibliowicz, J., Jeffery, W. R. and Rétaux, S. (2014). Enhanced prey capture skills in Astyanax cavefish larvae are independent from eye loss. Evodevo 5, 35.

Espinasa, L., Ornelas-García, C. P., Legendre, L., Rétaux, S., Best, A., Gamboa-Miranda, R., Espinosa-Pérez, H. and Sprouse, P. (2020). Discovery of Two New Astyanax Cavefish Localities Leads to Further Understanding of the Species Biogeography. Diversity 12, 368.

Fisher, R. A. (1930). The genetical theory of natural selection. 272,.

Grønskov, K., Ek, J. and Brondum-Nielsen, K. (2007). Oculocutaneous albinism. Orphanet J. Rare Dis. 2, 43.

Jeffery, W. R. (2001). Cavefish as a model system in evolutionary developmental biology. Dev. Biol. 231, 1–12.

Jeffery, W. R. (2005). Adaptive evolution of eye degeneration in the Mexican blind cavefish. J. Hered. 96, 185–196.

Jeffery, W. R., Strickler, A. G. and Yamamoto, Y. (2003). To see or not to see: evolution of eye degeneration in mexican blind cavefish. Integr. Comp. Biol. 43, 531–541.

Klaassen, H., Wang, Y., Adamski, K., Rohner, N. and Kowalko, J. E. (2018). CRISPR mutagenesis confirms the role of oca2 in melanin pigmentation in Astyanax mexicanus. Dev. Biol. 441, 313–318.

Kowalko, J. E., Rohner, N., Rompani, S. B., Peterson, B. K., Linden, T. A., Yoshizawa, M., Kay, E. H., Weber, J., Hoekstra, H. E., Jeffery, W. R., et al. (2013a). Loss of schooling behavior in cavefish through sight-dependent and sight-independent mechanisms. Curr. Biol. 23, 1874–1883.

Kowalko, J. E., Rohner, N., Linden, T. A., Rompani, S. B., Warren, W. C., Borowsky, R., Tabin, C. J., Jeffery, W. R. and Yoshizawa, M. (2013b). Convergence in feeding posture occurs through different genetic loci in independently evolved cave populations of Astyanax mexicanus. Proc. Natl. Acad. Sci. U. S. A. 110, 16933–16938.

Li, S., Li, H. and Takahata, T. (2022). Pigmented Long-Evans rats demonstrate better visual ability than albino Wistar rats in slow angles-descent forepaw grasping test. Neuroreport 33, 543–547.

Lin, J. Y. and Fisher, D. E. (2007). Melanocyte biology and skin pigmentation. Nature 445, 843–850.

Lloyd, E., Olive, C., Stahl, B. A., Jaggard, J. B., Amaral, P., Duboué, E. R. and Keene, A. C. (2018). Evolutionary shift towards lateral line dependent prey capture behavior in the blind Mexican cavefish. Dev. Biol. 441, 328–337.

Mackay, T. F. C. and Anholt, R. R. H. (2024). Pleiotropy, epistasis and the genetic architecture of quantitative traits. Nat. Rev. Genet.

Martin, A. and Orgogozo, V. (2013). The Loci of repeated evolution: a catalog of genetic hotspots of phenotypic variation. Evolution 67, 1235–1250.

Miranda-Gamboa, R., Espinasa, L., Verde-Ramírez, M. de L. A., Hernández-Lozano, J., Lacaille, J. L., Espinasa, M. and Ornelas-García, C. P. (2023). A new cave population of Astyanax mexicanus from Northern Sierra de El Abra, Tamaulipas, Mexico. SBC 45, 95–117.

Mitchell, R. W., Russell, W. H. and Elliott, W. R. (1977). Mexican eyeless Characin fishes, genus Astyanax: environment, distribution, and evolution.

Neuhauss, S. C. F. (2003). Behavioral genetic approaches to visual system development and function in zebrafish. J. Neurobiol. 54, 148–160.

Neuhauss, S. C., Biehlmaier, O., Seeliger, M. W., Das, T., Kohler, K., Harris, W. A. and Baier, H. (1999). Genetic disorders of vision revealed by a behavioral screen of 400 essential loci in zebrafish. J. Neurosci. 19, 8603–8615.

O’Gorman, M., Thakur, S., Imrie, G., Moran, R. L., Choy, S., Sifuentes-Romero, I., Bilandžija, H., Renner, K. J., Duboué, E., Rohner, N., et al. (2021). Pleiotropic function of the oca2 gene underlies the evolution of sleep loss and albinism in cavefish. Curr. Biol. 31, 3694-3701.e4.

Oliva, C., Hinz, N. K., Robinson, W., Barrett Thompson, A. M., Booth, J., Crisostomo, L. M., Zanineli, S., Tanner, M., Lloyd, E., O’Gorman, M., et al. (2022). Characterizing the genetic basis of trait evolution in the Mexican cavefish. Evol. Dev. 24, 131–144.

O’Quin, K. and McGaugh, S. E. (2016). Chapter 6 - Mapping the Genetic Basis of Troglomorphy in Astyanax: How Far We Have Come and Where Do We Go from Here? In Biology and Evolution of the Mexican Cavefish (ed. Keene, A.C.), Yoshizawa, M.), and McGaugh, S.E.), pp. 111–135. Academic Press.

Orr, H. A. (2000). Adaptation and the cost of complexity. Evolution 54, 13–20.

Otto, S. P. (2004). Two steps forward, one step back: the pleiotropic effects of favoured alleles. Proc. Biol. Sci. 271, 705–714.

Pantalia, M., Lin, Z., Tener, S. J., Qiao, B., Tang, G., Ulgherait, M., O’Connor, R., Delventhal, R., Volpi, J., Syed, S., et al. (2023). Drosophila mutants lacking the glial neurotransmitter-modifying enzyme Ebony exhibit low neurotransmitter levels and altered behavior. Sci. Rep. 13, 10411.

Patch, A., Paz, A., Holt, K. J., Duboué, E. R., Keene, A. C., Kowalko, J. E. and Fily, Y. (2022). Kinematic analysis of social interactions deconstructs the evolved loss of schooling behavior in cavefish. PLoS One 17, e0265894.

Paz, A., Holt, K. J., Clarke, A., Aviles, A., Abraham, B., Keene, A. C., Duboué, E. R., Fily, Y. and Kowalko, J. E. (2023). Changes in local interaction rules during ontogeny underlie the evolution of collective behavior. iScience 26, 107431.

Protas, M. E., Hersey, C., Kochanek, D., Zhou, Y., Wilkens, H., Jeffery, W. R., Zon, L. I., Borowsky, R. and Tabin, C. J. (2006). Genetic analysis of cavefish reveals molecular convergence in the evolution of albinism. Nat. Genet. 38, 107–111.

Protas, M., Conrad, M., Gross, J. B., Tabin, C. and Borowsky, R. (2007). Regressive evolution in the Mexican cave tetra, Astyanax mexicanus. Curr. Biol. 17, 452–454.

Protas, M., Tabansky, I., Conrad, M., Gross, J. B., Vidal, O., Tabin, C. J. and Borowsky, R. (2008). Multi-trait evolution in a cave fish, Astyanax mexicanus. Evol. Dev. 10, 196–209.

Reissmann, M. and Ludwig, A. (2013). Pleiotropic effects of coat colour-associated mutations in humans, mice and other mammals. Semin. Cell Dev. Biol. 24, 576–586.

Ren, J. Q., McCarthy, W. R., Zhang, H., Adolph, A. R. and Li, L. (2002). Behavioral visual responses of wild-type and hypopigmented zebrafish. Vision Res. 42, 293–299.

Rennison, D. J. and Peichel, C. L. (2022). Pleiotropy facilitates parallel adaptation in sticklebacks. Mol. Ecol. 31, 1476–1486.

Rodriguez-Morales, R., Gonzalez-Lerma, P., Yuiska, A., Han, J. H., Guerra, Y., Crisostomo, L., Keene, A. C., Duboue, E. R. and Kowalko, J. E. (2022). Convergence on reduced aggression through shared behavioral traits in multiple populations of Astyanax mexicanus. BMC Ecol Evol 22, 116.

Şadoğlu, P. (1957). A mendelian gene for albinism in natural cave fish. Experientia 13, 394– 394.

Sadoglu, P. and McKee, A. (1969). A second gene that affects eye and body color in Mexican blind cave fish. J. Hered. 60, 10–14.

Schindelin, J., Arganda-Carreras, I., Frise, E., Kaynig, V., Longair, M., Pietzsch, T., Preibisch, S., Rueden, C., Saalfeld, S., Schmid, B., et al. (2012). Fiji: an open-source platform for biological-image analysis. Nat. Methods 9, 676–682.

Sköld, H. N., Aspengren, S., Cheney, K. L. and Wallin, M. (2016). Fish Chromatophores--From Molecular Motors to Animal Behavior. Int. Rev. Cell Mol. Biol. 321, 171–219.

Slavík, O., Horký, P. and Wackermannová, M. (2016). How does agonistic behaviour differ in albino and pigmented fish? PeerJ 4, e1937.

Song, J., Yan, H. Y. and Popper, A. N. (1995). Damage and recovery of hair cells in fish canal (but not superficial) neuromasts after gentamicin exposure. Hear. Res. 91, 63–71.

Stern, D. L. (2013). The genetic causes of convergent evolution. Nat. Rev. Genet. 14, 751–764.

Van Trump, W. J., Coombs, S., Duncan, K. and McHenry, M. J. (2010). Gentamicin is ototoxic to all hair cells in the fish lateral line system. Hear. Res. 261, 42–50.

Warren, W. C., Boggs, T. E., Borowsky, R., Carlson, B. M., Ferrufino, E., Gross, J. B., Hillier, L., Hu, Z., Keene, A. C., Kenzior, A., et al. (2021). A chromosome-level genome of Astyanax mexicanus surface fish for comparing population-specific genetic differences contributing to trait evolution. Nat. Commun. 12, 1447.

Wilkens, H. (1988). Evolution and Genetics of Epigean and Cave Astyanax fasciatus (Characidae, Pisces). In Evolutionary Biology: Volume 23 (ed. Hecht, M.K.) and Wallace, B.), pp. 271–367. Boston, MA: Springer US.

Wright, S. (1929). Fisher’s Theory of Dominance. Am. Nat. 63, 274–279.

Wright, S. (1939). Statistical Genetics in Relation to Evolution. Hermann.

Yamamoto, Y., Byerly, M. S., Jackman, W. R. and Jeffery, W. R. (2009). Pleiotropic functions of embryonic sonic hedgehog expression link jaw and taste bud amplification with eye loss during cavefish evolution. Dev. Biol. 330, 200–211.

Yoshizawa, M., Goricki, S., Soares, D. and Jeffery, W. R. (2010). Evolution of a behavioral shift mediated by superficial neuromasts helps cavefish find food in darkness. Curr. Biol. 20, 1631–1636.

Yoshizawa, M., Yamamoto, Y., O’Quin, K. E. and Jeffery, W. R. (2012). Evolution of an adaptive behavior and its sensory receptors promotes eye regression in blind cavefish. BMC Biol. 10, 108.

